# Accelerated osteocytic citrate production in chronic kidney disease is associated with protection of the kidney

**DOI:** 10.1101/2025.10.08.681058

**Authors:** Jie Ren Gerald Har, Maggie Yun-Hsuan Hsu, Keith L. Saum, Akshdeep Singh, Ruchir Sriram, Dylan P. Kuennen, Emily K. Zhu, Subramaniam Pennathur, Maciek R. Antoniewicz, Lauren E. Surface

**Affiliations:** Department of Biologic and Material Sciences and Prosthodontics, University of Michigan School of Dentistry, Ann Arbor, MI, USA; Department of Chemical Engineering, University of Michigan, Ann Arbor, MI, USA; Division of Nephrology, Department of Internal Medicine, University of Michigan, Ann Arbor, Michigan, USA; Department of Molecular and Integrative Physiology, University of Michigan, Ann Arbor, Michigan, USA

**Keywords:** Chronic kidney disease, renal osteodystrophy, Bone metabolism, ^13^C-metabolic flux analysis, Citrate, SLC13A5, Nephrolithiasis

## Abstract

Patients with chronic kidney disease (CKD) face elevated fracture incidences, but mechanisms underlying CKD-related bone loss remain unclear. Using the adenine-induced chronic kidney injury (AdKI) murine model, we identified that AdKI induces dysregulated glucose metabolism in bones and kidneys via *in vivo* and *ex vivo* metabolic tracing. *Ex vivo* ^13^C-metabolic tracing of osteocyte-enriched femora revealed accelerated citrate production from [1,2-^13^C]-glucose and [U-^13^C]-glutamine in AdKI mice. These metabolic changes were observed together with increased circulating citrate and *Slc13a5* overexpression in bones from AdKI mice. Thus, to explore the role of citrate in AdKI, we utilized mice harboring a loss of function mutation in the citrate importer SLC13A5 (*Slc13a5*^*R337*/R337\**^). Mutant mice displayed elevated osteocytic citrate production, and elevated circulating citrate, without significantly worsened AdKI-related bone loss. Coincident with this, *Slc13a5*^*R337*/R337\**^ mutant mice were significantly protected from loss of kidney function with attenuated AdKI-induced nephrolithiasis. We also confirmed that *Slc13a5* is highly expressed in cortical bone compared to the kidney, suggesting the effect of the mutation is mediated by SLC13A5’s function outside the kidney. Altogether, this study finds that accelerated osteocytic citrate production in CKD is associated with protection of kidney function, and modulation of citrate handling may be a site for therapeutic intervention in CKD.

## Introduction

Chronic kidney disease (CKD) leads to systemic complications beyond the kidney, including bone loss and increased fracture risk^1–4^. This greatly reduces patient mobility and quality of life, and is a driver of mortality in CKD. In particular, patients with end-stage CKD face 2-3 times higher risk of hip fracture than their age- and sex-matched counterparts^4,5^. A high proportion of these patients present with high-turnover renal osteodystrophy (HT-ROD)^6–8^, characterized by bone loss concomitant with both increased bone formation and resorption rates^9,10^ and secondary hyperparathyroidism^6,8,11,12^. To manage bone loss in patients with CKD, there is an interest in investigating how skeletal metabolic phenotypes are disrupted by disease progression to modulate skeletal carbon metabolism as a novel strategy to improve bone health. This strategy aligns with recent studies on bone loss seen in mouse models of Type 1 diabetes^13^ or Type 2 diabetes^14^. In both cases, poor glycolysis was identified in the cortical bone of mice, and pharmacologic (metformin) or genetic strategies to improve skeletal glycolysis led to a reduction in diabetic bone loss. These studies provide a basis for research into the dysregulation of skeletal metabolism due to systemic diseases including diabetes^15,16^ or disuse osteoporosis^17^.

Investigating the dysregulation of skeletal metabolic fluxes in CKD-associated bone loss is particularly relevant, given previous investigations that discovered broad changes to skeletal metabolism in CKD. For example, a study utilizing Alport Syndrome mice (*Col4a3*^*−/−*^) and bone biopsy samples found reduced skeletal expression of *Hnf4a*, which encodes a regulator of glycolytic enzymes and lipid metabolism^18^, and identified that the loss of osteoblastic and osteocytic *Hnf4a* reduces osteogenesis *in vivo* and *in vitro*^19^, thus partially explaining bone loss in HT-ROD. In another study, bone loss in the adenine-induced CKD murine model was associated with mitochondrial dysfunction within osteocytes and can be partially rescued via mitochondrial antioxidants^20^. While these studies have described broad disruption to bioenergetic metabolism in the skeleton in complementary models of CKD, specific changes to metabolic fluxes or individual pathways are not yet recognized. To our present knowledge, high-resolution investigation into skeletal metabolic fluxes using stable-isotopic tracing technology has mostly been applied in studies on developing bone, skeletal stem cells^21–24^, and diabetic bone loss^13,14^.

Thus, identifying how specific metabolic fluxes in bone cells are altered in CKD, a disease with a very distinct effect on bones, will elucidate how bone metabolism may be driving disease progression. Given the complex crosstalk between the bone and kidney in CKD where bone-derived factors, including FGF23^25,26^ and sclerostin^27^, contribute to worsening of disease, understanding how bone metabolism is altered in CKD may also enable us to modulate the worsening function of the kidney itself.

To therefore understand how skeletal metabolic phenotypes are disrupted by CKD, we applied both *in vivo* and *ex vivo* ^13^C-metabolic tracing to osteocyte-enriched cortical bones from mice treated with the adenine-diet induced kidney injury (AdKI) model of CKD. We identified accelerated osteocytic citrate production in AdKI-associated bone loss, together with greater tibial expression of the SLC13 family of carboxylate transporters and elevated circulating citrate. We then utilized a mouse which harbors a loss-of-function mutation in the plasma membrane citrate transporter, SLC13A5^28^, to understand the role of citrate metabolism in AdKI. The mutation increased osteocytic citrate production, and had a significant protective effect on kidney function in AdKI mice. Altogether, these findings reveal a novel role of osteocytic citrate metabolism and the role of SLC13A5 in CKD progression on both bones and kidneys.

## Results

### Adenine diet-induced bone loss is observed together with systemic disruption of glucose metabolism

To investigate how CKD disrupts skeletal metabolism, we employed the established adenine diet-induced chronic kidney injury (AdKI) model in C57BL/6J mice. Previous studies have demonstrated in mice that this model of AdKI induces bone loss reminiscent of HT-ROD^9,10^. For our initial *in vivo* and *ex vivo* investigations, 8-week-old wildtype mice were induced for 4 weeks on the adenine diet, and then sacrificed, while age-matched controls were maintained over the same period on a control diet without adenine (Fig. 1A). μCT analysis of tibiae from these mice (Fig. 1B) revealed a significant loss in cortical bone (Fig. 1F-G) with weaker effects on trabecular bone (Fig. 1D-E), consistent with previous AdKI studies^9,20,29^. Gene set enrichment analysis using data from RNASeq of tibial cortices from control and AdKI mice (Table S1) revealed a pattern of increased bone resorption (Fig. 1J), as transcripts related to bone resorption were enriched in the upregulated set of genes (Fig. 1K), while the downregulated set was enriched for transcripts related to bone formation (Fig. 1L). Disruptions to circulating factors associated with CKD were also observed in the AdKI mice, including elevated circulating FGF23 and inorganic phosphate (Fig. 1H-I). Targeted GC-MS quantification of serum metabolites revealed systemic disruption of metabolites involved in bioenergetic processes. This includes an elevation in circulating glutamine, citrate, and glycine with decreased circulating glucose in AdKI mice (Fig. 1C, Supp. Fig 1). Disruptions to circulating glutamine, glycine, and glucose in CKD have been reported previously^30–32^. While reports differ on how citrate levels are altered in CKD, with many studies reporting decreased levels of citrate in patients with CKD^33,34^, and others reporting an association between higher circulating citate and decreasing kidney function^35–38^, several studies have suggested citrate may play a protective role in kidney injury and disease^39,40^.

**Figure 1:**
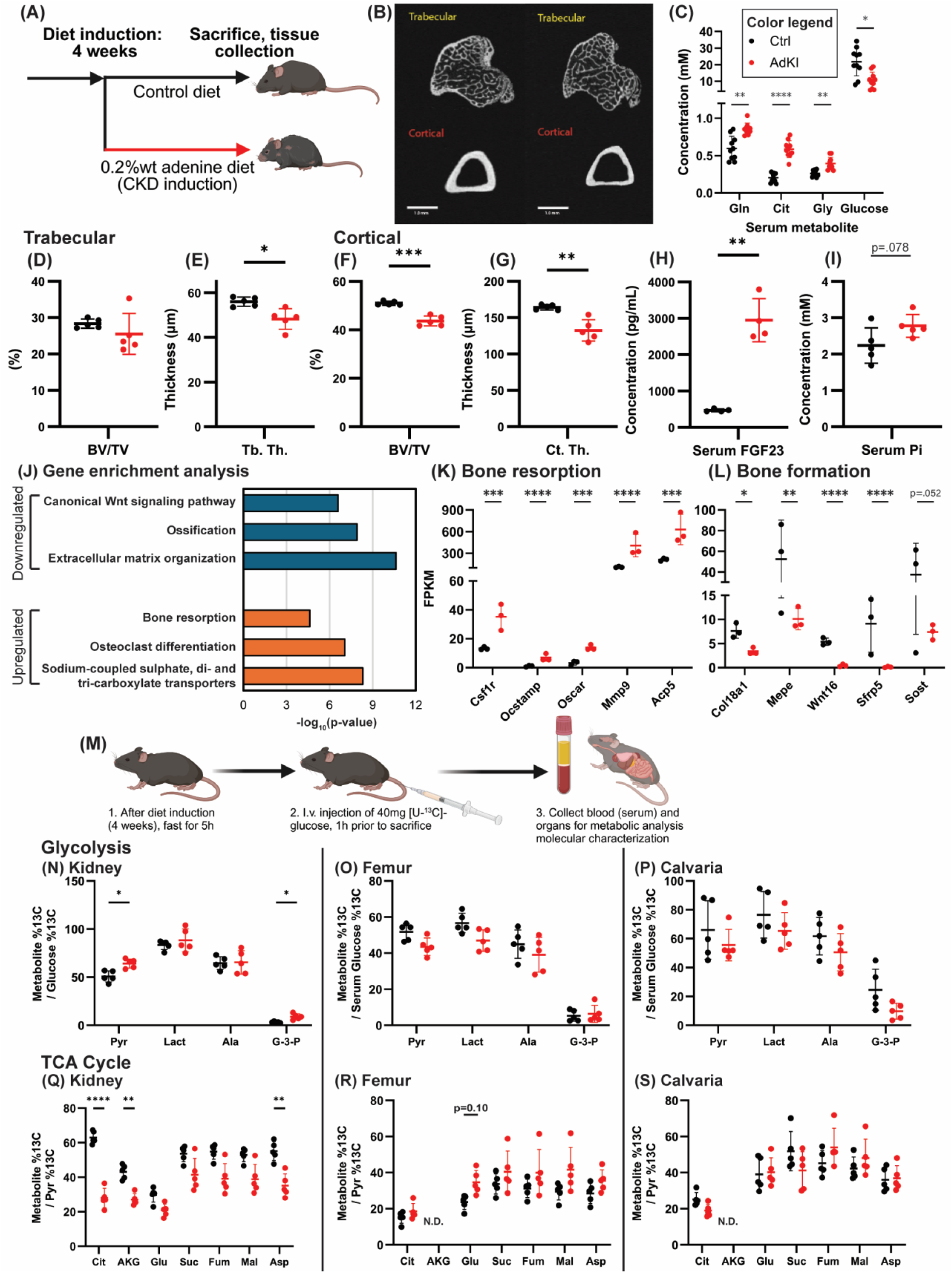
Bone loss in the adenine-diet induced chronic kidney injury (AdKI) murine model were observed together with disrupted *in vivo* glucose metabolism in bones and kidneys. (A) Diet schedule. (B) Representative tibia μCT images. (C) Selected circulating (serum) metabolites (n=10 per group). μCT quantification of (D-E) trabecular bone and (F-G) cortical bone (n=5 per group). Circulating (H) FGF23 (n=4 per group) and (I) inorganic phosphate (n=5 per group). (J) Gene enrichment analysis based on RNASeq of tibia cortices (n=3 per group). Gene expression levels, via RNASeq, of (K) selected genes involved in bone resorption and (L) bone formation, and statistical significance was determined via DEseq2. (M) Schematic of *in vivo* glucose tracing via intravenous injection of a bolus of [U-^13^C]-glucose. Relative glycolytic rate in (N) kidneys, (O) femora (flushed femur cortices), and (P) calvariae (biopsy punches of parietal bone); and relative TCA cycle activity in (Q) kidneys, (R) femora, and (S) calvariae. For *in vivo* metabolic tracing, n=5 per group. N.D. – AKG levels in femora and calvariae were below detection limit.

To examine how AdKI rewires organismal glucose metabolism, we performed *in vivo* stable isotope tracing in organs using [U-^13^C]-glucose^13,14^ in AdKI and control mice (Fig. 1M). In the kidneys, a kidney injury phenotype of accelerated glycolysis (Fig. 1N) and reduced glucose anaplerosis (Fig. 1Q) was observed in AdKI mice, consistent with recent studies^41–43^. In flushed femur cortices and calvarial punches, glycolytic and anaplerotic phenotypes were not significantly altered (Fig. 1O-P,R-S). Closer inspection revealed a trend of increased utilization of glucose in the synthesis of glutamate in femur cortices from AdKI compared to control mice (Fig. 1R). We confirmed that these *in vivo* glucose metabolic phenotypes in bones and kidneys were not primarily a result of hyperphosphatemia, by performing the same *in vivo* tracing of glucose metabolism on a separate group of mice induced on a 1.8% (w/w) high Pi diet (HPD) for 4 weeks along with controls (Supp. Fig. 2).

### Osteocyte-enriched bone from AdKI mice demonstrate accelerated ex vivo citrate production rates

To analyze in greater detail how metabolic phenotypes within the bone may be disrupted in AdKI bone loss, we performed *ex vivo* ^13^C-isotopic tracing on osteocyte-enriched femur cortices and calvarial punches using [1,2-^13^C]-glucose as a metabolic tracer (Fig. 2B). This *ex vivo* culture system (Fig. 2A) has been employed previously in studies examining osteocyte biology^44–46^. By quantifying the net consumption and production rates of metabolites of osteocyte enriched bone, we observed an increased osteocytic citrate production rate in bones from Ad-KI mice compared to control mice, particularly in calvariae where citrate production rate was 70% faster (p<.05) in AdKI calvariae (Fig. 2C-D).

**Figure 2:**
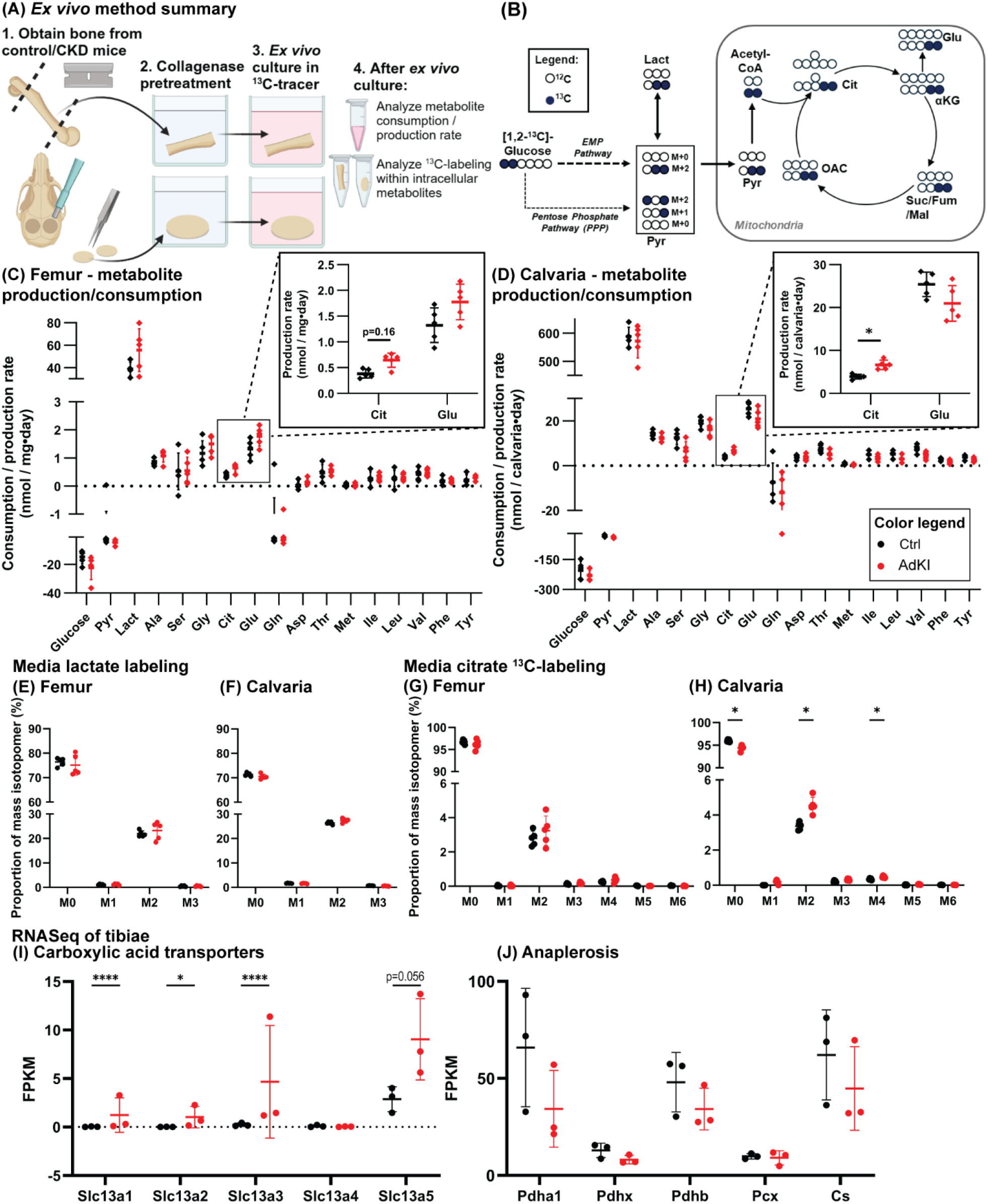
*Ex vivo* metabolic tracing using [1,2-^13^C]-glucose of osteocyte-enriched femora and calvariae revealed accelerated citrate production from glucose in osteocyte-enriched bones obtained from AdKI mice against that of controls. (A) Illustrated summary of *ex vivo* metabolic tracing – osteocyte-enriched femur cortices (flushed of marrow) and calvariae (biopsy punches of parietal bone) were obtained after repeated collagenase digestion. Following an overnight pre-culture in osteogenic culture media, *ex vivo* metabolic tracing in [1,2-^13^C]-glucose was performed on the osteocyte-enriched bones. (B) Schematic illustration of pertinent ^13^C-enriched metabolites. Net consumption and production rate of (C) osteocyte-enriched femora and (D) osteocyte-enriched calvariae. ^13^C-labeling of excreted lactate in the spent media of (E) femora and (F) calvariae cultures. ^13^C-labeling of excreted citrate in the spent media of (G) femora and (H) calvariae cultures (n=5 per group, for *ex vivo* metabolic analysis). Gene expression levels within tibia cortices of (I) genes encoding the SLC13 family of carboxylic acid transporters and (J) anaplerotic enzymes via RNASeq (n=3 per group, statistical significance was determined via Deseq2).

Relying on ^13^C-labeling data, we then investigated the nature of this accelerated citrate production in the osteocyte-enriched organ cultures. By analyzing the ^13^C-labeling patterns of extracellular citrate, we observed that calvaria punches from AdKI mice secreted significantly more citrate derived from [1,2-^13^C]-glucose over the course of the culture, suggesting the increased production of citrate by osteocyte-enriched bones is at least partially dependent on glucose. This is inferred from the greater proportion of both M2 and M4 citrate in the spent culture media of calvarial punches from AdKI mice compared to controls (Fig. 2H). In the *ex vivo* culture of femora cortices, while glucose was clearly employed in citrate production, we did not observe a significant increase in the proportion of glucose-derived citrate produced by AdKI femur cortices (Fig. 2G), perhaps due to incomplete removal of other cell types in our collagenase-digested femurs. In later experiments with an optimized digestion method (see Methods), AdKI significantly increases citrate production rate in osteocyte-enriched femora (Fig. 3A,E).

**Figure 3:**
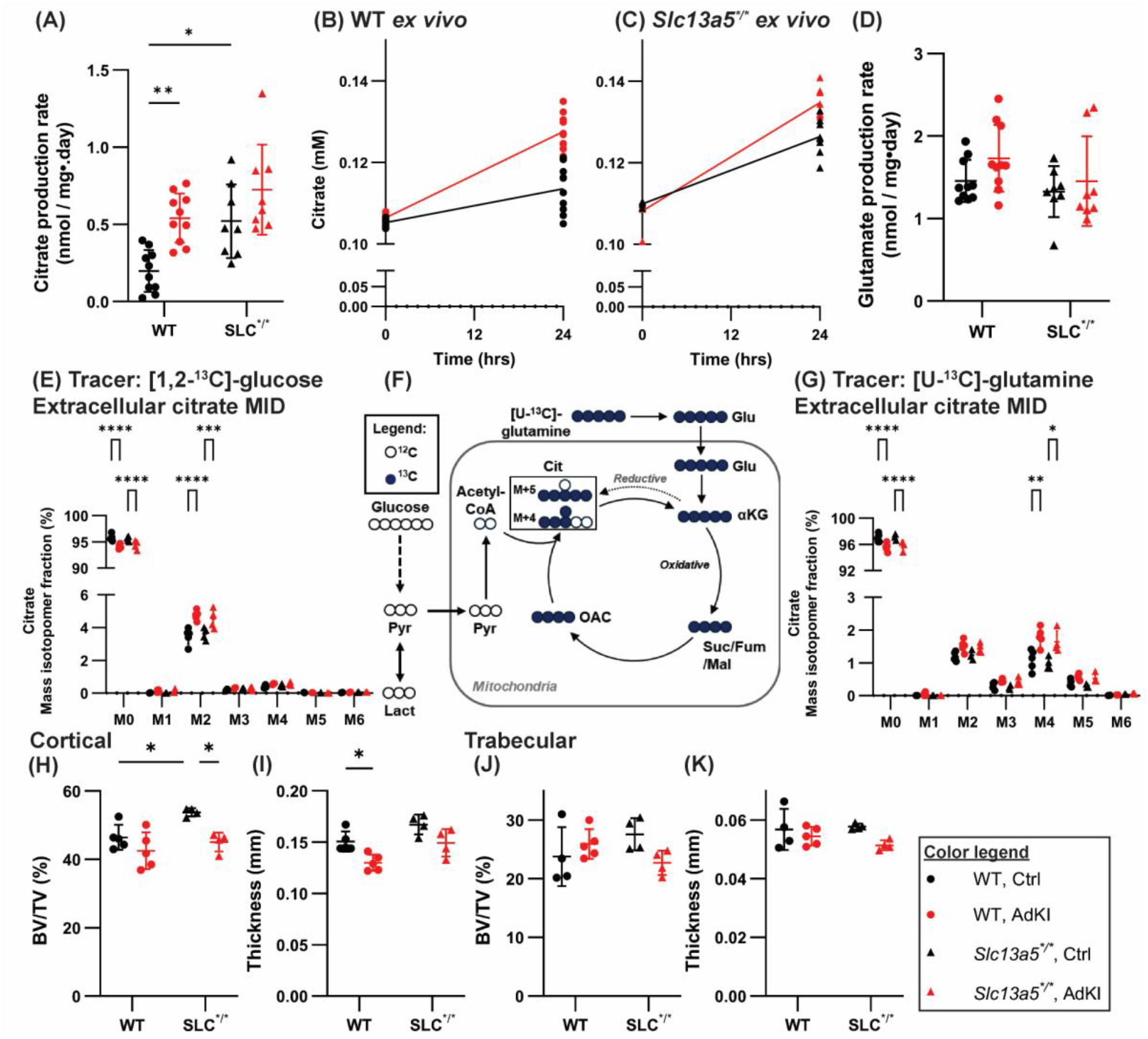
*Slc13a5*^*R337*/R337\**^ loss-of-function mutation increases osteocytic citrate production but does not significantly worsen AdKI-induced bone loss. Comparison of (A) *ex vivo* citrate production rate and (D) glutamate production rate. Concentration of citrate in culture media of osteocyte-enriched femora from (B) WT mice and (C) mutant mice (n=10 for WT and n=8 for *Slc13a5*^*\*/\**^ per group for *ex vivo* metabolic rates). (E) ^13^C-labeling of excreted citrate in the spent media of *ex vivo* Ocy-Fem cultured in [1,2-^13^C]-glucose. (F) Schematic illustration of pertinent ^13^C-enriched metabolites in [U-^13^C]-glutamine culture. (G) ^13^C-labeling of excreted citrate in the spent media of *ex vivo* Ocy-Fem cultured in [U-^13^C]-glutamine (n=5 for WT and n=4 for *Slc13a5*^*\*/\**^ per group for ^13^C-labeling analysis). Analysis of (H-I) cortical and (J-K) trabecular bone via μCT in 22 week-old male mice (n=5 for WT and n=4 for *Slc13a5*^*\*/\**^ per group for μCT analysis).

We also observed that accelerated glucose-dependent citrate production was a seemingly unique metabolic perturbation induced by AdKI. AdKI did not induce changes to either the production rate or the mass isotopomer distribution (MID) of other excreted metabolites (Table S2) that we analyzed including lactate (Fig. 2E-F) and glutamate (Fig. 2C-D, inset, Supp. Fig. 3D,J). Analysis of ^13^C-labeling in extracellular metabolites and metabolite production rates thus highlights increased osteocytic glucose-dependent citrate production rate as a unique metabolic disruption induced by AdKI.

Analysis of ^13^C-labeling of intracellular metabolites over 24 hours of *ex vivo* osteocyte-enriched organ culture (Supp. Fig. 4, Table S3) enabled inferences about whether intracellular metabolic pathways are altered due to AdKI. No significant differences were observed in the ^13^C-labeling patterns of intracellular glycolytic metabolites (pyruvate and lactate) over time, as is evident by the similarly increasing and then plateauing trends in M0 and M2 fragments within control and AdKI bone. As such, we infer that the ratio of glycolysis proceeding through either the pentose phosphate pathway (PPP) or the Embden-Meyerhof-Parnas (EMP) pathway remained unchanged, and the ^13^C-labeling of lactate and pyruvate in extracellular media (Fig. 2E-F, Supp. Fig. 3A-B,G-H) corroborates this.

Analysis of the ^13^C-labeling patterns in TCA cycle metabolites (Supp. Fig. 4C-F, I-L) revealed a trend of an increased proportion of intracellular M2 glutamate and M2 aspartate in the AdKI bones as compared to the controls. Particularly, the proportion of the M2 glutamate fragment within calvarial punches from AdKI mice was significantly greater than those from control mice (Supp. Fig. 4J). A similar trend, though statistically-insignificant, is also seen in aspartate extracted from calvaria punches (Supp. Fig. 4L) and glutamate and aspartate in femur cortices (Supp. Fig. 4D,F). This implied that a greater proportion of the glutamate and aspartate in AdKI osteocyte-enriched bone is derived from [1,2-^13^C]-glucose in the *ex vivo* culture media, echoing the trend of increased *in vivo* glucose-dependent synthesis of glutamate observed in femurs (Fig. 1R). Interestingly, despite this, there were no apparent changes in the ^13^C-labeling patterns of intracellular citrate and malate (Supp. Fig. 4C,E,I,K), and no significant change to the expression of genes coding for anaplerotic enzymes (Fig. 2J). These data together suggest that the increase in proportions of glucose-derived glutamate and aspartate is not due to an increase in glucose anaplerosis, but may reflect disrupted levels of glutamate and aspartate or nonessential amino acid metabolism.

### Loss-of-function mutation in Slc13a5 does not significantly worsen AdKI bone loss but ameliorates nephrolithiasis due to AdKI and improves kidney function

Taking these findings from extracellular and intracellular ^13^C-labeling together, we observed that accelerated osteocytic citrate production in AdKI bones is a product of glucose metabolism and not associated with broad changes in glycolytic metabolism or anaplerosis. This suggests glycolytic flux is not altered in these AdKI bones, but that mitochondrial dysfunction^20^ may be leading to a stalled TCA cycle resulting in an accumulation of citrate and increased glutamate synthesis. Since the skeleton is a storage of mineralized citrate^47,48^, the increase in osteocytic citrate production due to AdKI may have effects beyond the skeleton. We also observed, in addition to elevated the osteocytic citrate production rate, that AdKI bone loss was associated with an increase in circulating citrate (Fig. 1C) and overexpression of the SLC13 family of carboxylate transporters in the tibia cortices (Fig. 2I), including SLC13A5, a specialized plasma membrane citrate importer^28,49^. Given the prominence of disrupted citrate metabolism due to AdKI, we sought to analyze how exacerbating osteocytic citrate secretion via loss of the citrate importing function of SLC13A5 may impact AdKI disease progression in both the bones and kidneys. We hypothesized that loss of function of SLC13A5 would worsen the bone loss phenotype due to AdKI, based on a recent study that reported poorer femoral microarchitecture along with greater deposition of mineralized citrate in hindlimb long bones of *Slc13a5*^*-/-*^ knockout mice^49^. Separately, a Mendelian randomization study using Biobank data found that genetically proxied SLC13A5 inhibition is associated with better kidney function as illustrated by estimated glomerular filtration rate (eGFR) and lower blood urea nitrogen (BUN)^50^. Since SLC13A5 has been implicated in bone and kidney health, we then examined how loss of SLC13A5 function alters AdKI-related impaired bone and kidney function using the previously developed *Slc13a5*^*R337*/R337\**^ mutant mice^28^. This R337* mutation^28^, which results in a premature stop codon and organism-wide loss of citrate importer function, is modeled after the *SLC13A5*^*R333*/R333\**^ mutation found in patients with developmental epileptic encephalopathy and amelogenesis imperfecta.

We induced AdKI in 18-week-old male *Slc13a5*^*R337*/R337\**^ mutant and wildtype mice and analyzed bone and kidney function at 22 weeks, compared to mice maintained on the control diet. *Ex vivo* metabolic tracing of osteocyte-enriched femora from wildtype and mutant mice on the control diet revealed that the mutation resulted in a 2.7-fold increase (p<.05) in the net osteocytic citrate production rate (Fig. 3A), confirming that the loss-of-function mutation in SLC13A5 disrupts cellular citrate import and increases net citrate production. We also confirmed that osteocytic citrate production rate was increased by AdKI (Fig. 3A-C). This AdKI-induced increase in osteocytic citrate production was again unique to citrate – no significant change due to AdKI was observed in the glutamate production rate of femora, a closely-related TCA cycle metabolite and secreted during *ex vivo* culture (Fig. 3D). Using [1,2-^13^C]-glucose and [U-^13^C]-glutamine as metabolic tracers (Fig. 3E-G), we confirmed that the excreted citrate was actively produced from both glucose and glutamine, given the appearance of the M2 and M4 citrate fragments in the respective culture conditions. As previously, AdKI significantly increased the amount of M2 and M4 citrate produced from glucose and glutamine, respectively, in both wildtype and mutant bones (Fig. 3E,G, Table S4), consistent with earlier experiments that show active production of citrate from glucose (Fig. 2C-D, 2G-H).

To determine if increased osteocytic citrate production due to the *Slc13a5*^*R337*/R337\**^ mutation altered the extent of bone loss due to AdKI, μCT analysis of tibiae was performed (Supp. Fig. 5A-H). AdKI-induced bone loss in the cortical (Fig. 3H-I) and trabecular compartments (Fig. 3J-K) did not differ prominently between wildtype and mutant male mice, contrary to our hypothesis that AdKI bone loss may be worsened in the mutant mice. We then examined how this mutation alters AdKI-induced changes to circulating metabolites. Circulating citrate was significantly increased in both wildtype and mutant mice (Fig. 4A), confirming this systemic alteration in AdKI. Interestingly, circulating PTH, a key factor elevated in CKD, was significantly increased by AdKI in wildtype mice but not in mutant mice (Fig. 4B). Less severe hyperparathyroidism in AdKI mutant mice suggests that renal injury may be blunted in mutant mice. Furthermore, as elevated PTH (secondary hyperparathyroidism) is thought to contribute to CKD-related bone loss^51,52^, the expected AdKI bone loss in mutant mice may have been moderated by the reduced severity of hyperparathyroidism. Given the observation about circulating PTH and the role of citrate in kidney function and health^39,40,50^, we then analyzed how this *Slc13a5* mutation affects the progression of kidney disease in the AdKI mouse model.

**Figure 4:**
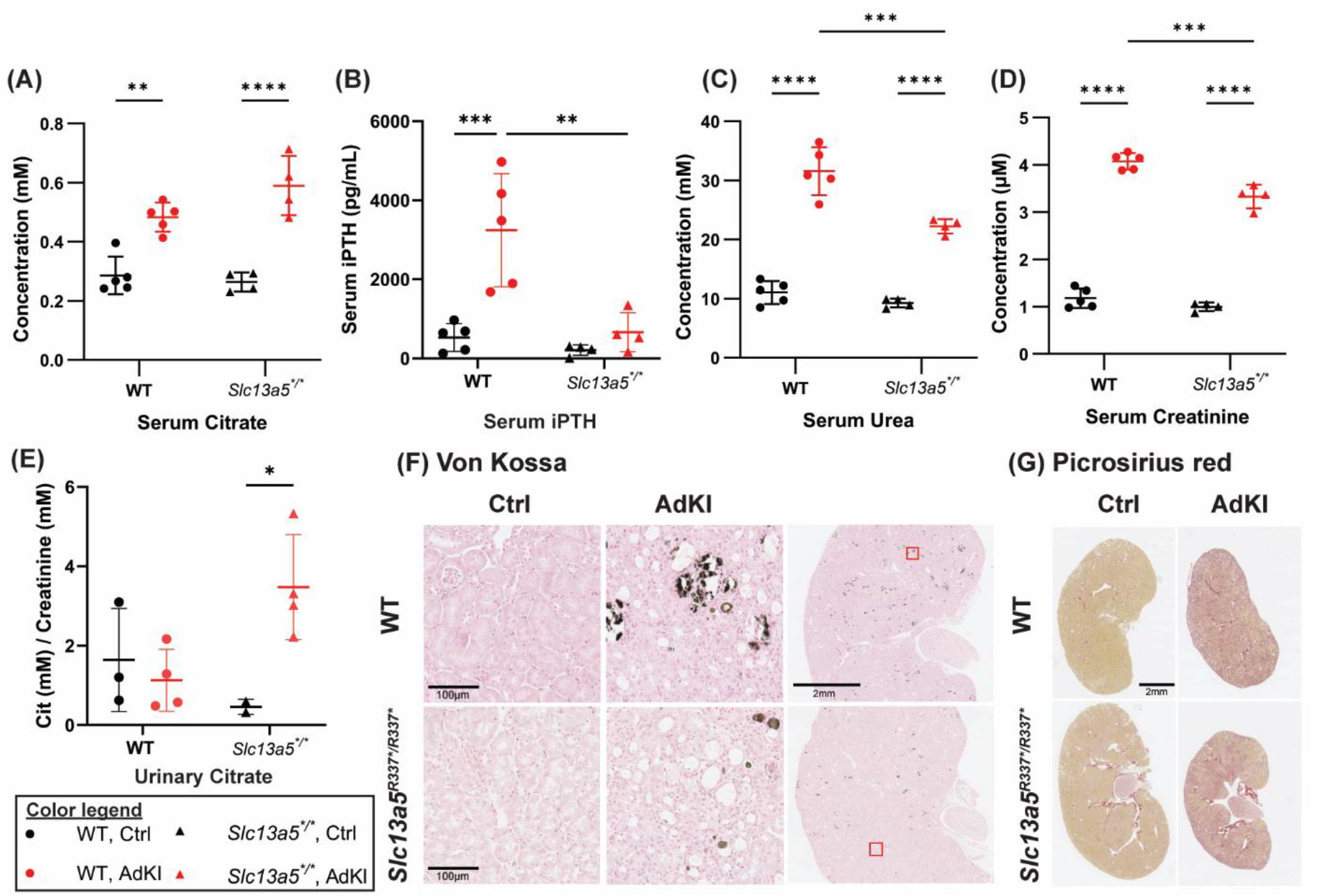
*Slc13a5*^*R337*/R337\**^ loss-of-function mutation significantly improves AdKI-induced decline in kidney function. Circulating (serum) levels of (A) citrate, (B) iPTH, (C) urea, and (D) creatinine (n=5 for WT and n=4 for *Slc13a5*^*\*/\**^ per group). (E) Urinary citrate clearance from spot urine of mice with sufficient urine (n=3 for WT Ctrl, n=2 for *Slc13a5*^*\*/\**^ Ctrl, and n=4 for WT and *Slc13a5*^*\*/\**^ AdKI). Representative images of stained kidney sections – (F) von Kossa and (G) picrosirius red.

The AdKI-induced increase in markers of renal dysfunction – serum urea and creatinine (Fig. 4C-D) – was also reduced in the AdKI-induced *Slc13a5*^*R337*/R337\**^ mutant mice, further suggesting less severe kidney disease progression due to loss of SLC13A5 function. This trend was comparable in both male and female mice (Supp. Fig. 7D-E). We then performed histological analysis of kidney samples to visualize the extent of kidney injury. *Slc13a5*^*R337*/R337\**^ mutant mice had significantly reduced nephrolithiasis following AdKI compared to wildtype mice, as von Kossa-stained kidney sections revealed a reduction in the extent and number of calcified tubules (Fig. 4F, Supp. Fig. 8A). This observation corresponds with elevated urinary citrate in the mutant AdKI mice (Fig. 4E), consistent with a role of urinary citrate in the suppression of kidney stone formation^53^. Picrosirius red-stained sections also indicate that renal fibrosis was lower in *Slc13a5*^*R337*/R337\**^ mutant mice compared to wildtype (Fig. 4G, Supp. Fig. 8B). Together, inhibition of SLC13A5 function blunted the severity of AdKI, evidenced by reduced nephrolithiasis and moderated elevation of CKD markers (PTH, urea, creatinine) in the *Slc13a5*^*R337*/R337\**^ mutant mice compared to wildtype mice.

## Discussion and Limitations

In this report, we describe how bone loss resulting from adenine diet-induced kidney injury associates with disruption in skeletal metabolism through the application of both *in vivo* and *ex vivo* ^13^C-metabolic tracing. Notably, *ex vivo* ^13^C-metabolic tracing revealed an increase in osteocytic citrate production due to AdKI. By employing ^13^C-labeled glucose and glutamine as metabolic tracers, we also confirmed that accelerated citrate production is a metabolically active process where osteocytes convert both substrates into citrate, and not just an artefact of matrix-bound citrate being released into the culture media. We note how these high-resolution metabolic tracing experiments complement previous findings on how kidney disease impacts metabolic phenotypes in the skeleton. For example, an earlier study identified decreased mitophagy in osteocytes of AdKI mice and a lowered rate of oxidative phosphorylation in primary osteoblasts from these mice^20^, suggesting a reduction in the TCA cycle activity of the bones of AdKI mice. Indeed, our experiments reveal greater osteocytic citrate production, likely corresponding to poorer citrate catabolism in TCA cycle, as well as faster enrichment of glucose-derived M2 glutamate and aspartate in AdKI bones (Supp. Fig. 4D,F,J,L). Altogether our data and others suggest AdKI disrupts TCA cycle activity in osteocyte-enriched cortical bone.

Accelerated osteocytic citrate production together with overexpression of *Slc13a5* (Fig. 2I), perhaps in response to the elevated extracellular citrate, and increased circulating citrate (Fig. 1C) establishes that AdKI disrupts skeletal citrate metabolism. As the skeleton serves as a major reservoir of organismal citrate^47,48^, further investigation into the role of skeletal citrate metabolism in AdKI is warranted. In this work, using mice harboring the *Slc13a5*^*R337*/R337\**^ mutation in the citrate importer SLC13A5, we found that exacerbating osteocytic citrate production did not significantly worsen bone loss due to AdKI. This contrasts with our expectations – given that young, 6-week-old male and female *Slc13a5*^*-/-*^ mice show poorer femoral microarchitecture compared with their controls^49,54^, we had anticipated that increased osteocytic citrate production due to the mutation would further worsen bone loss in mutant mice. One possible explanation is that the *Slc13a5*^*R337*/R337\**^ mutation confers protection to the kidneys, as is evident through lower circulating urea and creatinine levels, significant reduction in hyperparathyroidism, and markedly reduced nephrolithiasis in AdKI mutant mice (Fig. 4). These findings corroborate the recent GWAS study which identified a role of *SLC13A5* inhibition in improving eGFR and reducing BUN, and supports the potential of targeting the SLC13 family of transporters and citrate metabolism in metabolic and kidney diseases^50,55^. This protection of kidney function in the *Slc13a5*^*R337*/R337\**^ mutant mice may have moderated the systemic dysregulations due to AdKI, including less severe hyperparathyroidism in mutants (Fig. 4B), which would in turn have tempered the AdKI bone loss seen in the mutant mice.

A straightforward explanation for the protective effect of the *Slc13a5*^*R337*/R337\**^ mutation on kidney function may be that the loss of SLC13A5 function in kidneys of mutant mice directly increases urinary citrate in AdKI (Fig. 4E), thus protecting against stone formation^53^ and tubular injury due to AdKI. Yet, present studies on organismal-wide expression of *Slc13a5* complicate this inference, as *Slc13a5* expression is high in cortical bones but not the kidneys^49,56^. Our qRT-PCR data confirms higher *Slc13a5* expression in the bones than the kidneys (Supp. Fig. 9). As such, it is also possible that in the global loss of function SLC13A5 mutant mice, elevated citrate production from outside the kidney, including from the bone, the major storage site of citrate in the body, contributes to the protection of kidney function. These data suggest a potentially underappreciated role of osteocytic citrate as a metabolic regulator or marker of CKD progression, adding to the growing awareness of bone-derived factors in the progression of CKD^26,27,57^. However, future investigations to elucidate the role of osteocytic citrate metabolism using osteocyte-specific *Slc13a5* knockout mice will be critical to understanding the specific impact of skeletal citrate metabolism on CKD pathogenesis.

While we observe decreased adenine-induced kidney injury in female *Slc13a5*^*R337*/R337\**^ mutant mice (Supp. Fig. 7) as well as male mice, we acknowledge that this study has been conducted primarily using male mice and the AdKI model of CKD. It is well-understood that CKD progression induces differential increase in fracture risk in male and female patients^1,2^. Recent metabolomic analyses in mice show that the cortical bone metabolome is distinct between male and female mice^58^, and that AdKI in these mice produces distinct sexually dimorphic disruptions^9,59^. As such, future studies should investigate whether this is reflected in sexually dimorphic disruptions in skeletal metabolism in CKD. Additionally, further investigation using other non-nephrolithiasis models of kidney disease^60^ such as the 5/6-nephrectomy model or drug-induced glomerulosclerosis would confirm if this disruption in osteocytic citrate metabolism is consistent across models of bone loss in kidney disease, and whether loss of SLC13A5 function dampens kidney disease progression in a range of CKD models.

In summary, we identified accelerated osteocytic citrate production together with elevated circulating citrate and cortical bone expression of *Slc13a5* in the adenine-induced kidney injury mouse model of high-turnover bone loss. In investigating the role of the citrate transporter SLC13A5 on AdKI associated disruptions to bone and kidney function using the *Slc13a5*^*R337*/R337\**^ mutant mice, we observed that the mutation confers a protective effect of the kidneys in AdKI. This study provides a basis for studies into the role of citrate in CKD progression, especially in the case of nephrolithiasis as targeting citrate metabolism can be therapeutically beneficial for patients with CKD without compromising skeletal health.

## Resource Availability

### Lead contact

Further information and requests for resources and reagents should be directed to and will be fulfilled by the lead contact, Lauren Surface.

### Materials, data and code availability

This study did not generate new reagents or materials. RNASeq data set will be uploaded when GEO is operational (at time of submission, GEO is unavailable). ^13^C-metabolic tracing data are available as stated in Supplemental Tables. Images of scanned slides and other additional information required to reanalyze the data reported in this paper is available from the lead contact upon request.

## Acknowledgments

This study was supported by the University of Michigan Biological Sciences Scholar Program. L.E.S. is supported by NIH R00AR073903 (NIH/NIAMS) and R56DE033668 (NIH/NIDCR). We thank the Michigan Integrative Musculoskeletal Health Center (MIMHC) Structure, Composition and Histology Core for assistance and advice on μCT analysis of long bones (supported by NIH/NCRR S10RR026475-01) and the Orthopaedic Research Laboratories Histology Core for processing, sectioning and histological staining of kidneys. We thank Dr. Jan C-C. Hu for generously providing the *Slc13a5*^*R337*/R337\**^ mutant mice used in this study. Illustrations in Figure 1 and 2 were created in BioRender. Surface, L. (2025) https://BioRender.com/1kym2v4.

## Author Contributions

Conceptualization—J.R.G.H. and L.E.S.; Investigation and data collection—J.R.G.H., Y-H.M.H., R.S., A.S., D.P.K., E.Z., K.L.S., and L.E.S.; writing, original draft—J.R.G.H. and L.E.S.; writing, review & editing—J.R.G.H., L.E.S., K.L.S., D.P.K., S.P., M.R.A.; funding acquisition, L.E.S.; resources, L.E.S., S.P., and M.R.A.; supervision, L.E.S.

## Declaration of Interests

The authors declare no competing interests.

## Declaration of Generative AI and AI-Assisted Technologies

The authors did not use any generative AI or AI-assisted technology in the preparation of this work.

## Supplemental Information

**Document S1: Figures S1–S9**

**Supplemental Tables – refer to Mendeley Data: 10.17632/9sr7pb6c76.1**.

Table S1: Differentially expressed genes in tibia cortices of control and AdKI mice (n=3 per group), analyzed via RNASeq and the Deseq2 algorithm.

Table S2: Extracellular ^13^C-labeling of metabolites within spent media of osteocyte-enriched femora (Ocy-Fem) and osteocyte-enriched calvariae (Ocy-Cal) following 24h of *ex vivo* culture in [1,2-^13^C]-glucose (n=5 per group).

Table S3: Intracellular ^13^C-labeling of metabolites extracted from osteocyte-enriched femora (Ocy-Fem) and osteocyte-enriched calvariae (Ocy-Cal) following 24h of *ex vivo* culture in [1,2-^13^C]-glucose (n=5 per group). Note: At the start (time 0h), it is assumed that M0 fragments predominate (100%) and no ^13^C-enrichment has taken place yet. Data at 9h and 24h are extracted from *ex vivo* cultured osteocyte-enriched bone organs.

Table S4: Extracellular ^13^C-labeling of metabolites within spent media of osteocyte-enriched femora (Ocy-Fem) following 24h of *ex vivo* culture in [1,2-^13^C]-glucose or [U-^13^C]-glutamine (n=5 for control and AdKI WT mice; n=4 for control and AdKI *Slc13a5*^*R337*/R337\**^ mice).

## Experimental Model

### Animals and disease model

To investigate adenine diet-induced chronic kidney injury (AdKI) in mice, mice were induced on a 0.2% (w/w) adenine diet (Inotiv TD.160020) over 4 weeks. To investigate if hyperphosphatemia exerts an influence on *in vivo* metabolism (Supp. Fig. 2), mice were induced on a high phosphate (1.8%) diet over 4 weeks, and age-matched controls for these were maintained on a control phosphate diet (0.6%) over the same period.

In earlier experiments (Fig. 1 and 2, Supp. Fig. 1-4), 8-week-old wildtype C57BL/6J male mice (Jackson Laboratories) were induced on the adenine diet, control diet (Inotiv TD.150303), or high phosphate diet or phosphate control diet for 4 weeks. At the end of diet induction (12 weeks of age), mice were euthanized via ketamine/xylazine anesthesia and exsanguination followed by cervical dislocation.

The mutant *Slc13a5*^*R337*/R337\**^ mice were developed^28^ and generously provided by Dr. Jan C-C. Hu for this study. 18-week-old male *Slc13a5*^*R337*/R337\**^ male mice, 18-week-old wildtype male mice (Jackson Laboratories) (Fig. 3-4, Supp. Fig. 5,8), 13-week-old *Slc13a5*^*R337*/R337\**^ female mice, 13-week-old wildtype female mice (Supp. Fig. 6-7) were randomly separated and then induced on the adenine diet and control diet for 4 weeks. At the end of diet induction (males 22 weeks of age, females 17 weeks of age), mice were euthanized via exsanguination under isoflurane followed by secondary cervical dislocation. Sera and organs were collected for metabolic analysis and other molecular characterization.

## Method

### Materials

[U-^13^C]-glucose powder was purchased from Cambridge Isotope Laboratories. Complete MEMα was purchased from Gibco. Depleted MEMα (MEMα with nucleosides, less glucose, pyruvate, aspartate, asparagine, glutamate, glutamine, and ascorbic acid) for ex vivo metabolic tracing was purchased from Nucleus Biologics. Fetal bovine serum (FBS, FetalClone III) was purchased from Cytiva. All other common reagents are summarized in the resource table below.

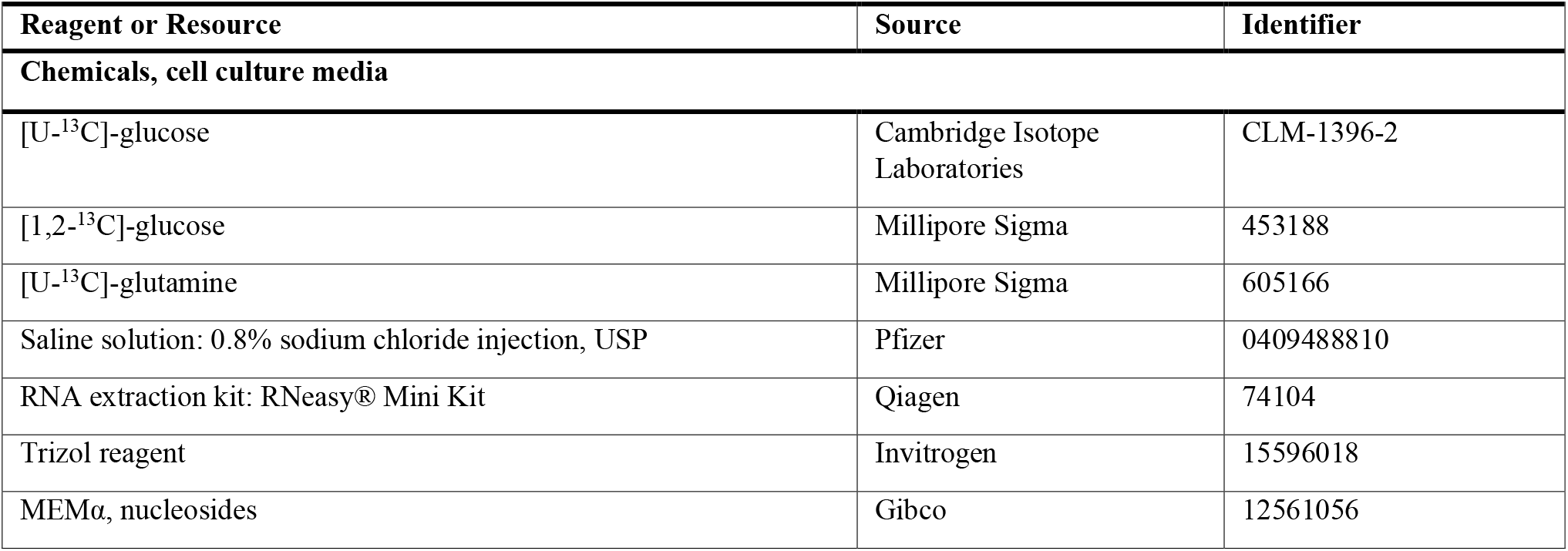
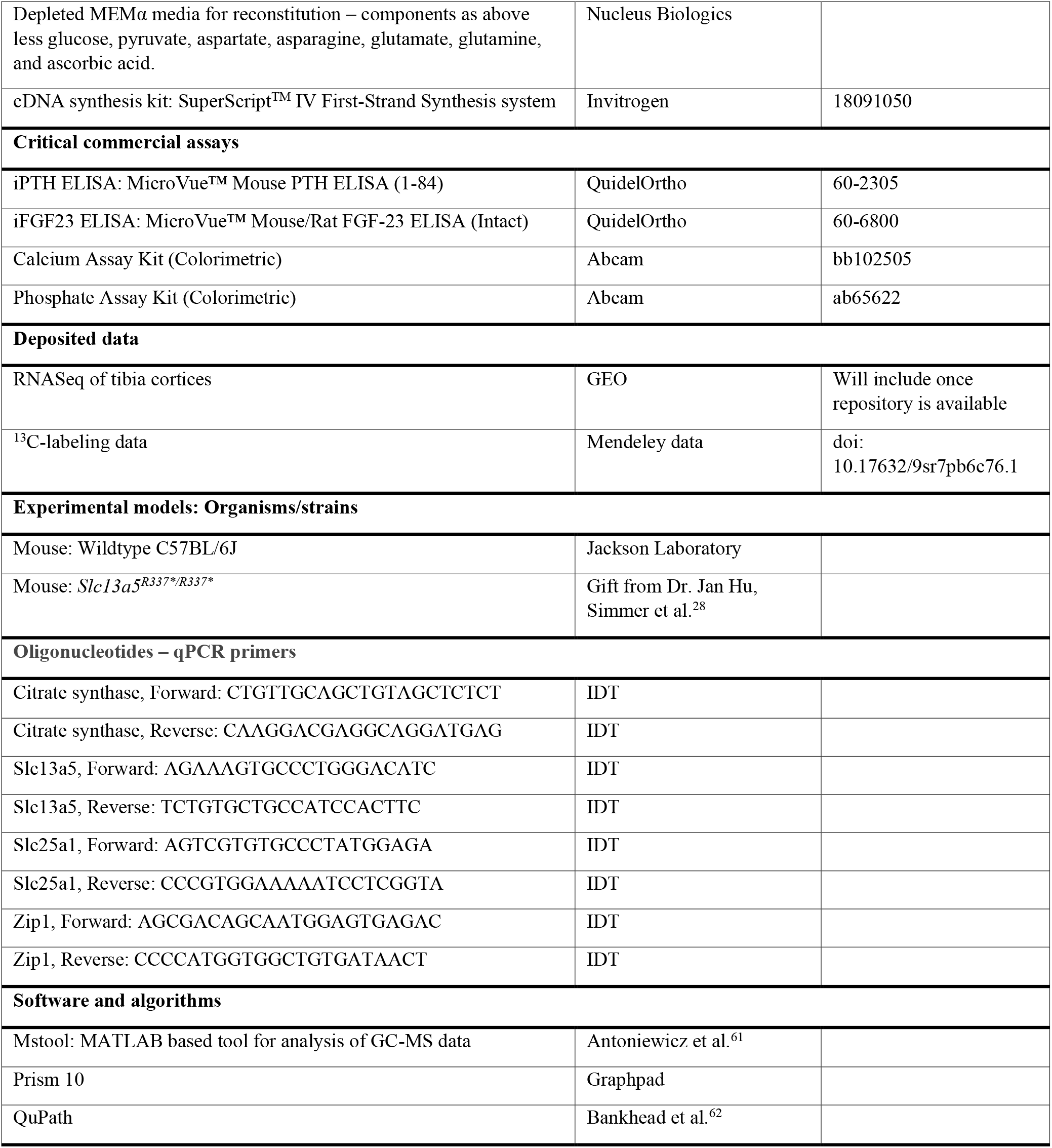

### *In vivo* ^13^C-tracing for metabolic analysis

At the end of the 4 weeks of diet induction, each mouse was fasted for 5h and then injected with 20 mg of [U-13C]-glucose dissolved in sterile saline. [U-13C]-glucose solution was prepared at 400 mg/mL. The mouse was then left in a cage unrestrained for 1h and then euthanized. Serum, and organs were collected quickly and flash frozen. For metabolic analysis of bone tissue, the ends of femora were cut off and marrow was flushed with PBS prior to freezing; and calvariae, including parietal and interparietal bone, were cleaned of connective tissue prior to freezing.

### *Ex vivo* ^13^C-tracing for metabolic analysis in the bone

At the end of diet induction, following euthanasia, flushed diaphyses of femora and calvaria punches (from the parietal bone) were obtained quickly and placed in warm, complete MEMα media + 10% FBS prior collagenase treatment. In the earlier, exploratory experiments (8-week-old male mice, AdKI against control), flushed diaphyses of the femora and calvaria were digested sequentially with Type I collagenase (3 mg/mL) in HBSS with 0.1% BSA at 37°C to remove surface lining cells. Each round of digestion was performed for 30min and the supernatant was discarded after each round. A total of 4 rounds of digestion was performed.

Thereafter, the osteocyte-enriched femora and calvaria were rinsed thoroughly in HBSS and conditioned in complete MEMα overnight. Bone samples were then rinsed thoroughly with HBSS and cultured in reconstituted MEMα + 10% FBS, in which unlabeled glucose was replaced with [1,2-^13^C]-glucose for *ex vivo* metabolic tracing.

In later experiments (18-week-old male mice, wildtype and *Slc13a5*^*R337*/R337\**^, AdKI against control), flushed diaphyses of the femora were digested sequentially with modifications to improve the osteocytic fraction^45,63^. First, 3 rounds of digestion with Type I collagenase (3 mg/mL) in complete MEMα media with 1% BSA (MEMα + collagenase) at 37°C for 30min was performed, and the supernatant was discarded after each round. Following this, the flushed diaphyses were digested in 5 mM EDTA in HBSS with 1% BSA (EDTA in HBSS) at 37°C for 30min, and the supernatant was discarded. Following this, alternating digestion in MEMα + collagenase and EDTA in HBSS was performed for another 3 rounds. After a total of 9 sequential digests, the osteocyte-enriched femora and calvaria were rinsed thoroughly in HBSS and conditioned in complete MEMα overnight. Bone samples were then rinsed thoroughly with HBSS and cultured in reconstituted MEMα + 10% FBS, in which unlabeled glucose was replaced with [1,2-^13^C]-glucose or unlabeled glutamine was replaced with [U-^13^C]-glutamine for *ex vivo* metabolic tracing.

### Serum

Blood was left to clot on ice for 1-2h, and then serum was collected by centrifugation of clot blood at 2000g for 15min. Serum was frozen prior to subsequent analysis. Assay kits for the quantification of FGF23, PTH, phosphate, and calcium are listed in the Key Resources Table. Absorbance values were read using the Tecan Spark multimode microplate reader. Concentration of small metabolites in serum were measured via GC-MS, except for creatinine which was measured via LC-MS^64^.

### Metabolite extraction and GC-MS analysis of ^13^C-labeling in metabolites

The method employed was adapted from previous studies^14,65,66^. Organs collected and flash frozen following metabolic tracing experiments were cryo-pulverized with a tissue crusher and quickly placed into 1.8 mL of 1:1:1 chloroform/methanol/water mixture (extraction solvent) in a glass tube. Extraction of metabolites from serum was similarly performed by placing 30 μL of serum into 1.8 mL of extraction solvent. The suspension was then vortexed vigorously for 30s.

The aqueous (top) layer, containing extracted polar metabolites, was then carefully aspirated and transferred into a clean microcentrifuge tube and lyophilized at 4°C for downstream analysis via GC-MS.

Lyophilized metabolites extracted from organs or sera were then derivatized and ^13^C-labeling was measured using GC-MS as previously performed^65,66^. Analysis of mass isotopomer data was performed using Mstool. The reader is referred to Tables S2-S4 for analyzed mass isotopomer distribution data.

### Quantification of serum metabolites

Briefly, for quantification of metabolites via GC-MS, an equal volume of serum was mixed with an equal volume of [U-^13^C]-labeled standard solution with known concentrations of amino acids and metabolites of interest. This mixture was then placed into 1 mL of extraction solvent and the aqueous fraction was then lyophilized for downstream derivatization and analysis via GC-MS to determine of concentrations of each metabolite present in serum or culture media based on the ratio of unlabeled (sample) to ^13^C-labeled (standard) metabolite fragments^65,66^. Quantification of creatinine via LC-MS was performed as described^64^.

### MicroCT

Fixed and dehydrated tibiae were placed in a 19 mm diameter specimen holder and scanned using a µCT system (µCT100 Scanco Medical, Bassersdorf, Switzerland). Scan settings were: voxel size 12 µm, 70 kVp, 114 µA, 0.5 mm AL filter, and integration time 500 ms. Analysis was performed using XQuartz (Version 2.8.5), and a fixed global threshold of 18% (180 on a grayscale of 0 – 1000) for trabecular bone and 28% (280 on a grayscale of 0 – 100 was used to segment bone from non-bone. A 0.5 mm region of trabecular bone was analyzed immediately below the growth plate and a 0.5 mm cortical slice was analyzed around the midpoint of the tibiae.

### RNASeq

RNA from pulverized tibiae diaphyses flushed of marrow was extracted with RNeasy kit (Qiagen). For RNASeq and bioinformatics analysis (Novogene), sequencing was performed on the NovaSeq X Plus Series (PE150) platform, with 6G raw data per sample. Bioinformatics analysis was performed on QC, cleaned reads by mapping reads using Hisat v2.0.5 to the Mus musculus genome, and the DESeq2 algorithm was employed for differential expression analysis (Table S1). Gene enrichment analysis of significantly upregulated or downregulated genes was performed with Metascape^67^ (http://metascape.org).

### qRT-PCR

For qRT-PCR, RNA from pulverized tibiae diaphyses flushed of marrow or pulverized kidney halves were extracted using the Trizol reagent. RNA were redissolved in appropriate amount of RNAase-free water (about 20-50 μL). cDNA was then synthesized using the SuperScript™ IV First-Strand Synthesis system from 1 μg of RNA. qRT-PCR was performed with the above primers on the QuantStudio™ 3 System.

### Kidney histology

Kidneys were fixed overnight in formalin and stored in 70% ethanol at 4°C prior to further histological analysis by the University of Michigan Orthopaedic Research Laboratory Histology Core facility. To process kidneys for histological staining, tissues were dehydrated in a graded series of ethanol washes, processed, and paraffin-embedded and then sectioned at 5 μm thickness. Kidney sections mounted on glass slides were stained with von Kossa (counterstain: nuclear fast red), Picrosirius Red, and Masson’s Trichrome. Quantification of kidney calcification or fibrosis was performed using QuPath v0.5.1.

## Quantification and Statistical Analysis

Unless stated otherwise, comparison between two groups were performed using unpaired t-tests. Where multiple comparisons were made, t-tests were performed with the Bonferonni-Dunn correction. Where comparison between multiple groups were made, two-way ANOVA was employed and the Bonferonni correction was performed to correct for multiple comparisons. Legend for corrected p-values: *p<0.05, **p<0.01, ***p<0.005, ****p<0.001. P-values of interest greater than 0.05 are indicated above comparisons. Number of replicates in figure legends indicate biological replicates, unless stated otherwise. Statistical analysis and graphing was performed using Graphpad Prism 10.

## Notes

https://data.mendeley.com/

### Competing Interest Statement

The authors have declared no competing interest.

