## Supplemental Figures for "Accelerated osteocytic citrate production in chronic kidney disease is associated with protection of the kidney"

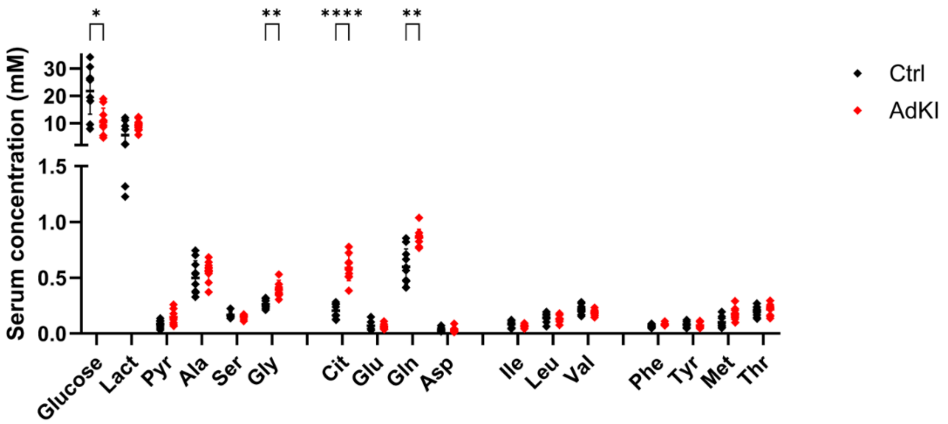


**Supplemental Figure 1:** Panel of serum metabolites from young adult male mice, measured via GC-MS (n=10). *p<.05, **p<.01, ***p<.005, ****p<.001 by t-tests with Bonferroni correction for multiple comparisons.

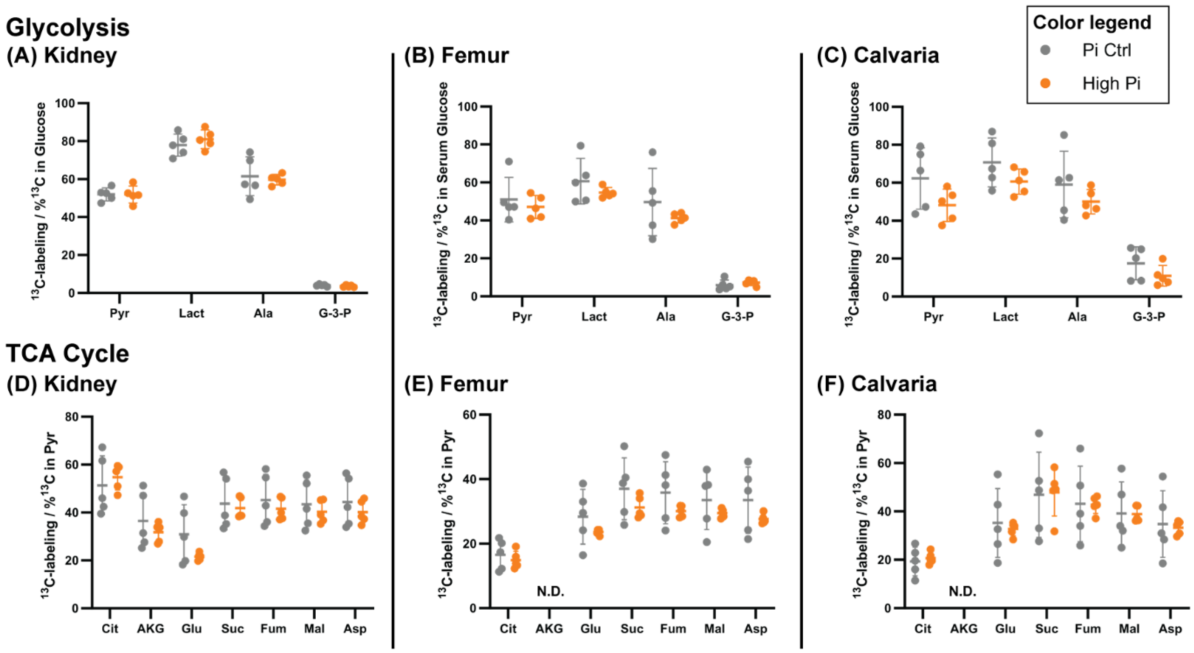


**Supplemental Figure 2:** *In vivo* glucose tracing via intravenous injection of a bolus of [U-$^{13}$C]-glucose in mice fed with 1.8% Pi diet (High Pi) against controls (Pi Ctrl, 0.6% Pi) (n=5 per group). Relative glycolytic rate in (A) kidneys, (B) femora (flushed femur cortices), and (C) calvariae; and relative TCA cycle activity in (D) kidneys, (E) femora, and (F) calvariae. N.D. no data collected for AKG in femora and calvariae due to low ion abundance.

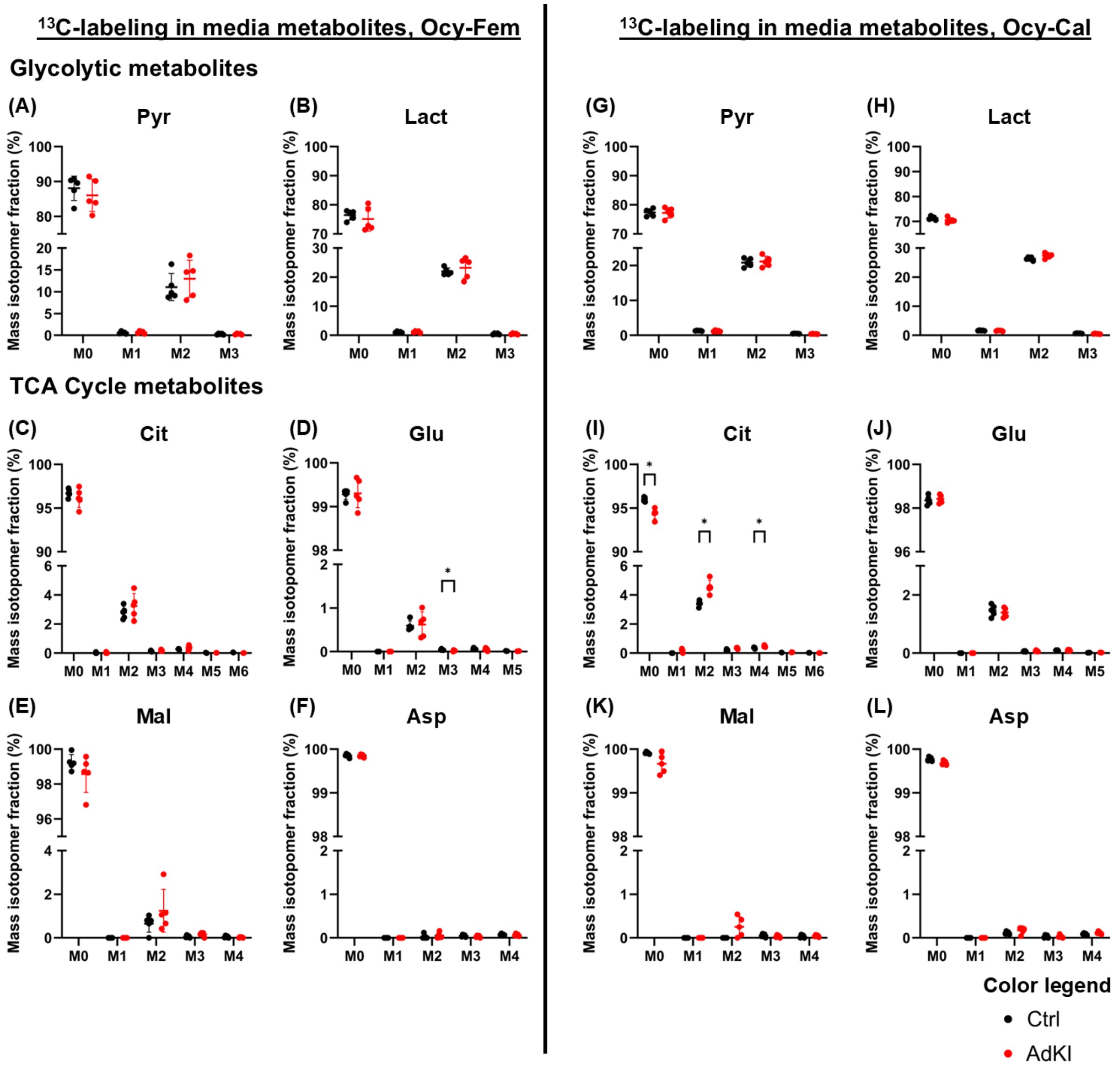


**Supplemental Figure 3:** $^{13}$C-labeling of metabolites in the culture media following 24h of *ex vivo* osteocyte-enriched bone organ culture with [1,2-$^{13}$C]-glucose as a metabolic tracer. Within the spent media of osteocyte-enriched femora (Ocy Fem), $^{13}$C-labeling of: (A) pyruvate; (B) lactate; (C) citrate; (D) glutamate; (E) malate; and (F) aspartate. Within the spent media of osteocyte-enriched calvariae (Ocy-Cal), $^{13}$C-labeling of: (G) pyruvate; (H) lactate; (I) citrate; (J) glutamate; (K) malate; and (L) aspartate. n=5 per group, *p<.05 via t-tests with Bonferroni correction for multiple comparisons.

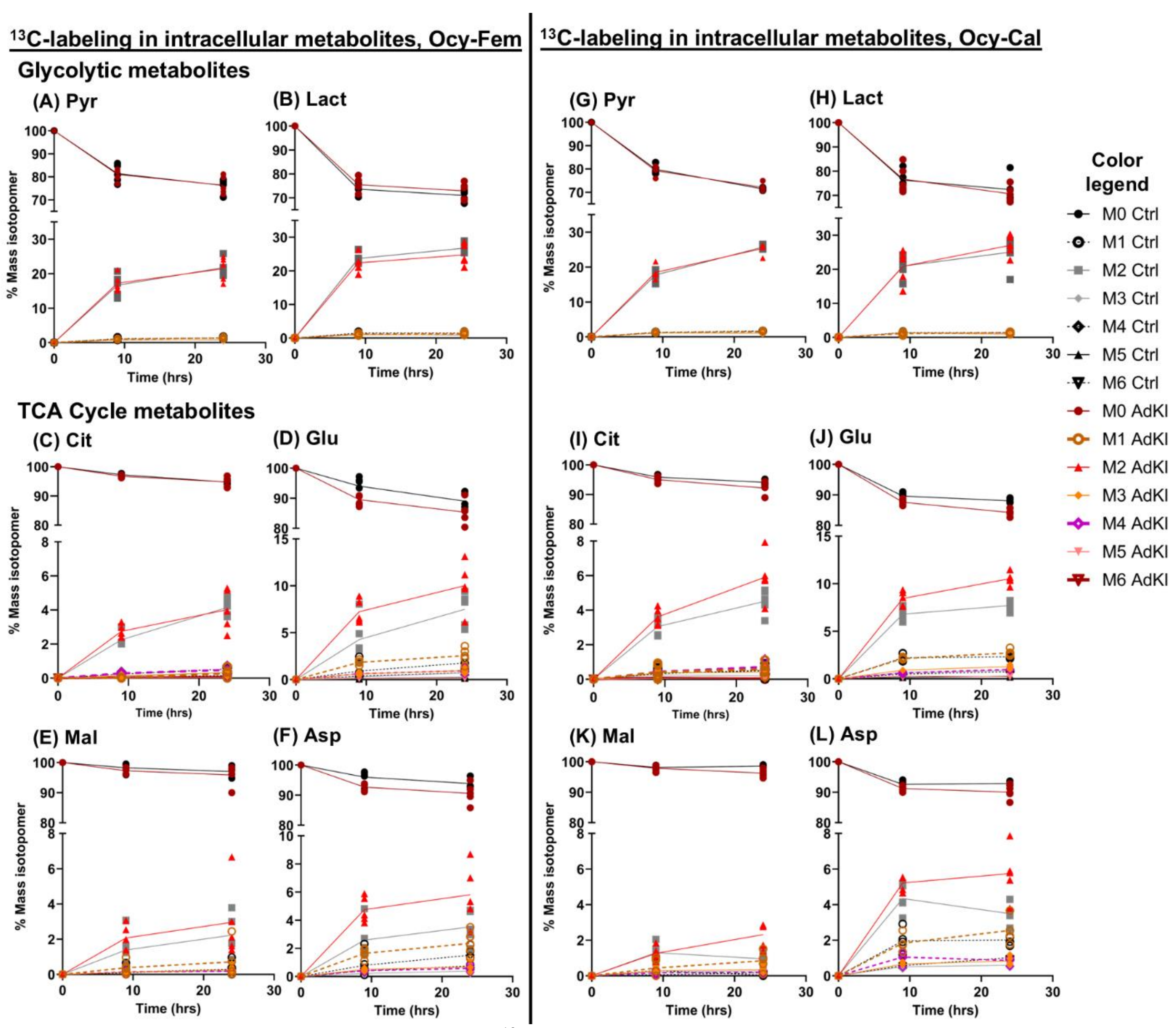


**Supplemental Figure 4:** Intracellular metabolite $^{13}C$-labeling patterns extracted from osteocyte-enriched femora (Ocy Fem) and osteocyte-enriched calvariae (Ocy-Cal) of young adult male mice (AdKI and control) cultured ex vivo with [1,2-$^{13}C$]-glucose as a metabolic tracer, at 9h and 24h. Mass isotopomer distributions of metabolites extracted from Ocy-Fem: (A) pyruvate; (B) lactate; (C) citrate; (D) glutamate; (E) malate; and (F) aspartate. Mass isotopomer distributions of metabolites extracted from Ocy-Cal: (G) pyruvate; (H) lactate; (I) citrate; (J) glutamate; (K) malate; and (L) aspartate.

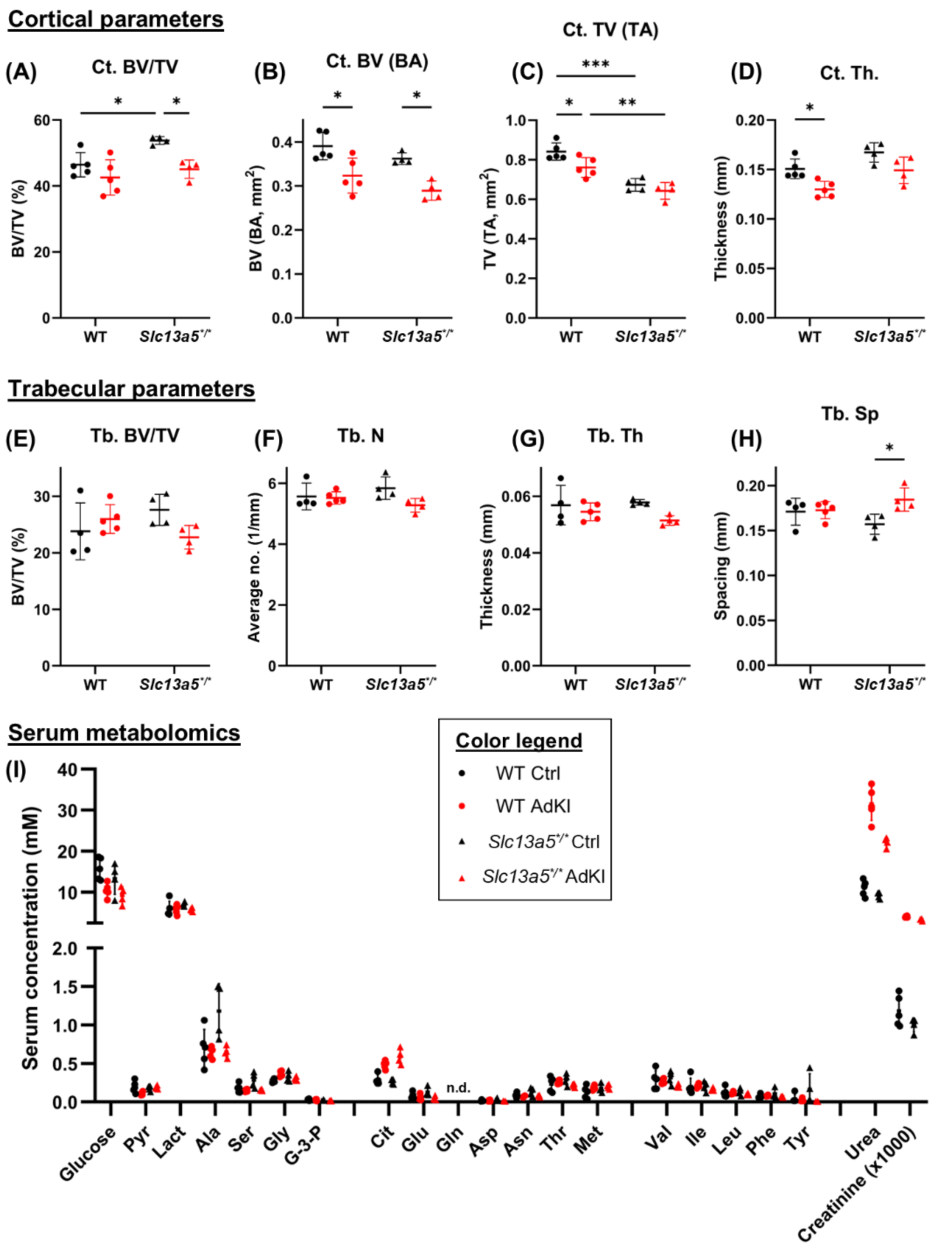


**Supplemental Figure 5:** µCT characterization of tibiae and serum metabolomics of male, adult (22-wk-old) *Slc13a5*$^{R337*/R337*}$ mice. Cortical parameters: (A) BV/TV; (B) bone volume; (C) tissue volume; and (D) cortical thickness. Trabecular parameters: (E) BV/TV; (F) trabecular number; (G) trabecular thickness; and (H) trabecular spacing. (I) Panel of serum metabolites from adult male mice (n=5 for WT and n=4 for *Slc13a5*$^{R337*/R337*}$). *p<.05, **p<.01, ***p<.005, ****p<.001 by ANOVA and post-hoc t-tests with Bonferroni correction for multiple comparisons.

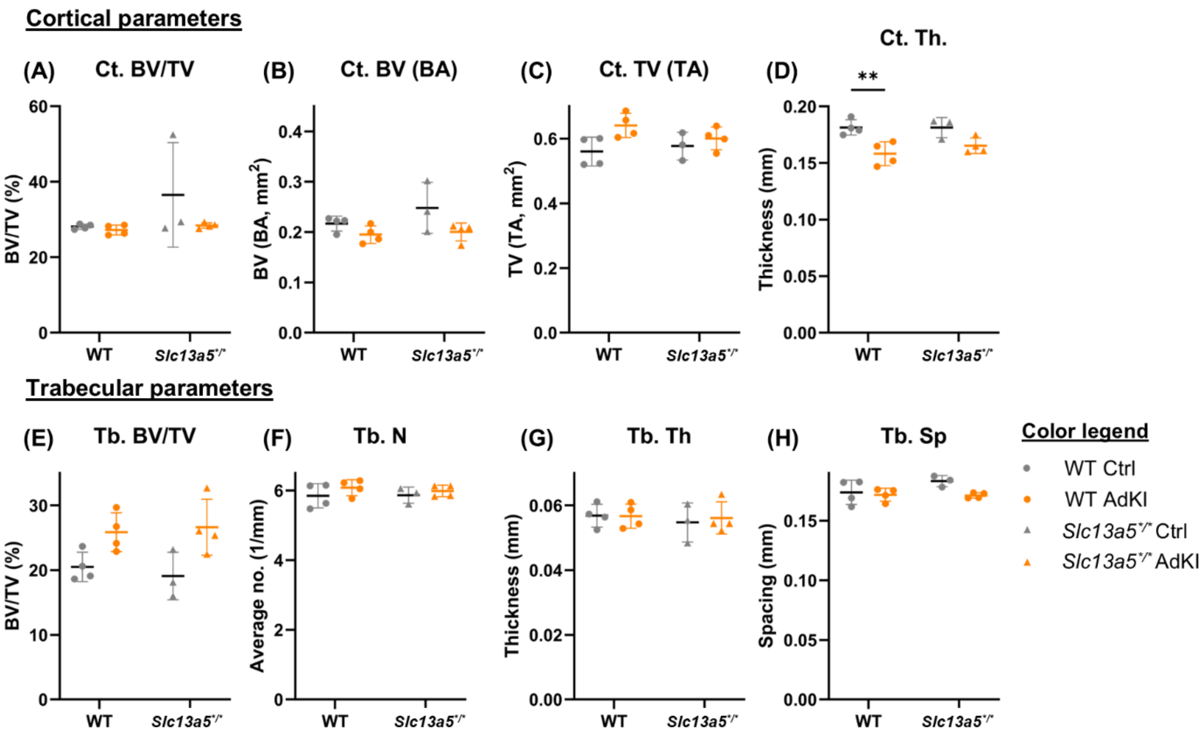


**Supplemental Figure 6:** µCT characterization of tibiae and serum metabolomics of female, adult (17-wk-old) *$Slc13a5^{R337*/R337*}$* mice. Cortical parameters: (A) BV/TV; (B) bone volume; (C) tissue volume; and (D) cortical thickness. Trabecular parameters: (E) BV/TV; (F) trabecular number; (G) trabecular thickness; and (H) trabecular spacing (n=4 for WT and *$Slc13a5^{R337*/R337*}$* AdKI, n=3 for *$Slc13a5^{R337*/R337*}$* Ctrl). *p<.05, **p<.01, ***p<.005, ****p<.001 by ANOVA and post-hoc t-tests with Bonferroni correction for multiple comparisons.

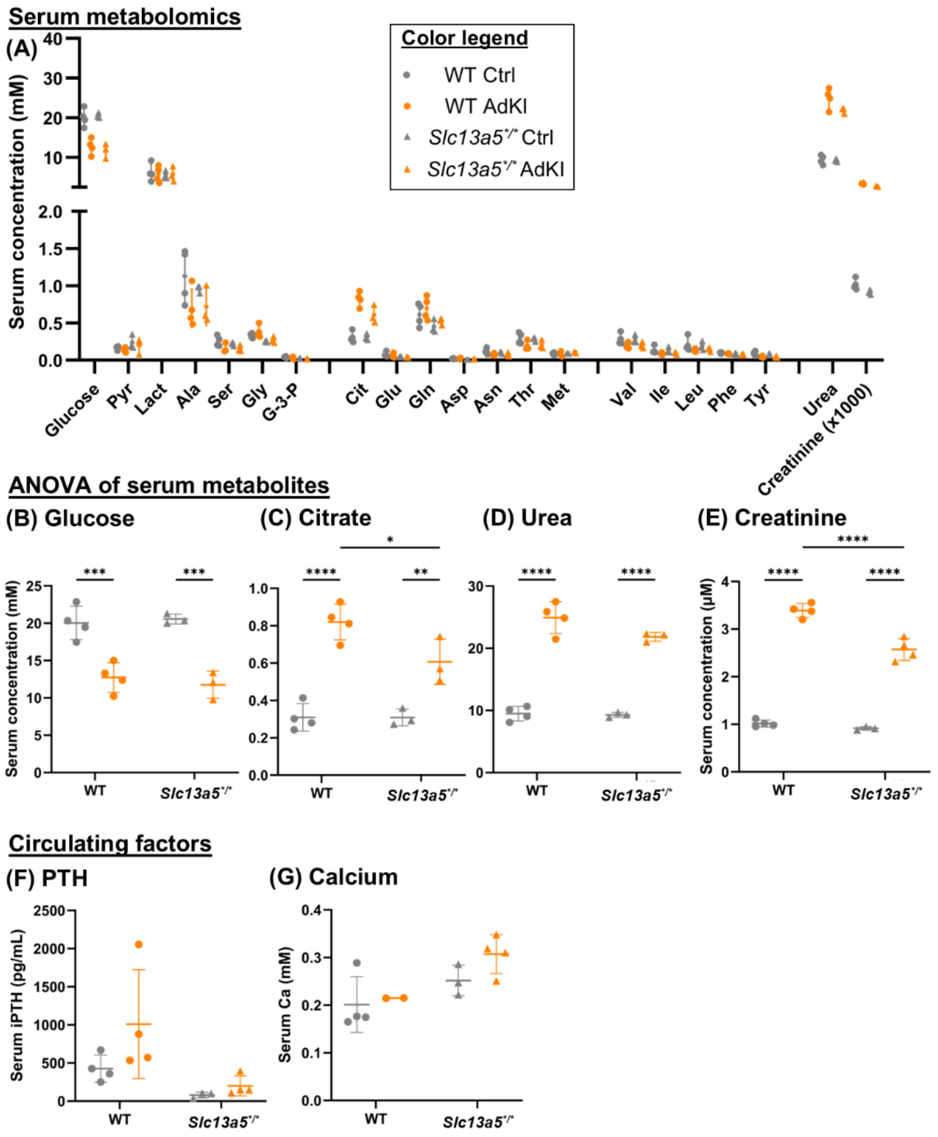


**Supplemental Figure 7:** (A) Panel of serum metabolites from adult female mice, measured via GC-MS (n=4 for WT and *Slc13a5*$^{R337*/R337*}$ AdKI, n=3 for *Slc13a5*$^{R337*/R337*}$ Ctrl). Statistical analysis (ANOVA) of serum (B) glucose, (C) citrate, (D) urea, and (E) creatinine. Circulating factors measured via ELISA and colorimetric assay respectively: (F) PTH and (G) ionic calcium. *p<.05, **p<.01, ***p<.005, ****p<.001 by ANOVA and post-hoc t-tests with Bonferroni correction for multiple comparisons.

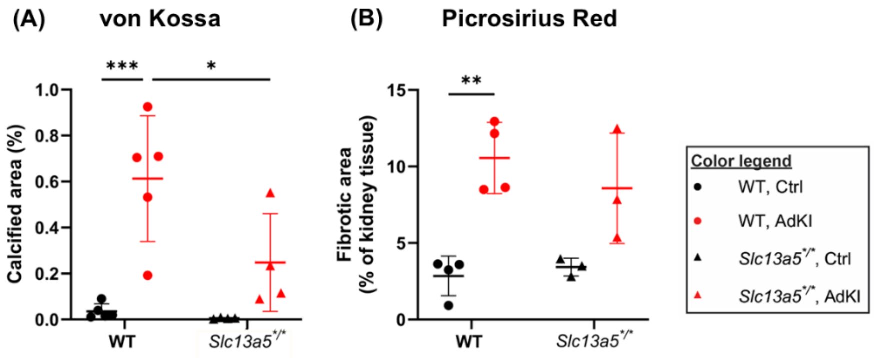


**Supplemental Figure 8:** Quantification of histologically-stained kidney sections. (A) Extent of calcification through von Kossa staining, and extent of fibrosis through (B) picrosirius red.

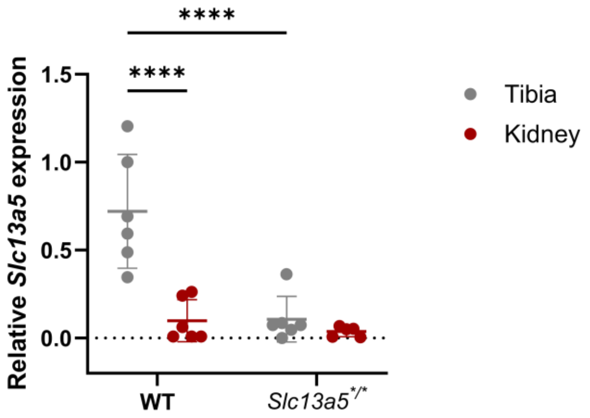


**Supplemental Figure 9:** qPCR analysis of Slc13a5 expression in the cortical bone of tibiae (flushed of marrow) and in kidneys (bulk).
